# Relict groups of spiny frogs indicate Late Paleogene-Early Neogene trans-Tibet dispersal of thermophile faunal elements

**DOI:** 10.1101/2021.02.18.430751

**Authors:** Sylvia Hofmann, Daniel Jablonski, Spartak Litvinchuk, Rafaqat Masroor, Joachim Schmidt

**Affiliations:** Centre of Taxonomy and Evolutionary Research, Zoological Research Museum Alexander Koenig, Bonn, Germany; Department of Zoology, Comenius University in Bratislava, Bratislava, Slovakia; Institute of Cytology, Russian Academy of Sciences, St. Petersburg, Russia; Zoological Sciences Division, Pakistan Museum of Natural History, Islamabad, Pakistan; Institute of Biosciences, General and Systematic Zoology, University of Rostock, Rostock, Germany

## Abstract

**Background:** The Himalaya-Tibet orogen (HTO) presents an outstanding geologically active formation that contributed to, and fostered, modern Asian biodiversity. However, our concepts of the historical biogeography of its biota are far from conclusive, as are uplift scenarios for the different parts of the HTO. Here, we revisited our previously published data set of the tribe Paini extending it with sequence data from the most western Himalayan spiny frogs *Allopaa* and *Chrysopaa* and using them as an indirect indicator for the paleoecological development of Tibet.

**Methods:** We obtained sequence data of two mitochondrial loci (16S rRNA, COI) and one nuclear marker (Rag1) from *Allopaa* samples from Kashmir Himalaya as well as *Chrysopaa* sequence data from the Hindu Kush available from GenBank to complement our previous data set. A Maximum likelihood and dated Bayesian gene tree were generated based on the concatenated data set. To resolve the inconsistent placement of *Allopaa,* we performed different topology tests.

**Results:** Consistent with previous results, the Southeast Asian genus *Quasipaa* is sister to all other spiny frogs. The results further reveal a basal placement of *Chrysopaa* relative to *Allopaa* and *Nanorana* with an estimated age of *ca.* 26 Mya. Based on the topology tests, the phylogenetic position of *Allopaa* as a sister clade to *Chaparana* seems to be most likely, resulting in a paraphyletic genus *Nanorana* and a separation from the latter clade around 20 Mya. Both, the placements of *Chrysopaa* and *Allopaa* support the presence of basal Paini lineages in the far north western part of the HTO, which is diametrically opposite end of the HTO with respect to the ancestral area of spiny frogs in Southeast Asia. These striking distributional patterns can be most parsimoniously explained by trans-Tibet dispersal during the late Oligocene (subtropical *Chrysopaa*) respectively early Miocene (warm temperate *Allopaa).* Within spiny frogs, only members of the monophyletic *Nanorana+Paa* clade are adapted to the colder temperate climates, indicating that high-altitude environments did not dominate in the HTO before *ca.* 15 Mya. Our results are consistent with fossil records suggesting that large parts of Tibet were characterized by subtropical to warm temperate climates at least until the early Miocene.

## Introduction

The uplift of the modern Himalaya-Tibet orogen (HTO) was one of the most extensive geological events during the Cenozoic. Today’s dimension of the HTO is thought to exert profound influences on the regional and global climate, and, consequently, on Asian biodiversity. Thus, understanding the evolution and knowing the past topography of the HTO is critical for exploring its paleoenvironments and historical biogeography (Kutzbach et al. 1989; Molnar et al. 2010; Raymo & Ruddiman 1992; Zhang et al. 2018). However, various lines of geoscientific evidence have suggested – partly substantially – different uplift scenarios for the respective parts of the HTO (reviewed in Spicer et al. 2020). These scenarios range from the idea of a simple monolithic rising of Tibet purely due to crustal thickening or lithosphere modification (e.g., Wang et al. 2014; Zhao & Morgan 1985), over different models of a fractional, stepwise development (e.g., Tapponnier et al. 2001), to the concept of a high ‘proto-Tibetan Plateau’ (Mulch & Chamberlain 2006; Wang et al. 2014). Linked to these varying conceptions are uncertainties in timing, quantity (elevational increase) and sequence pattern of the HTO uplift. While several geoscientific studies present evidence for a high elevated Tibetan Plateau (TP) as early as the Eocene or even earlier (e.g., Kapp et al. 2007; Murphy et al. 1997; Tapponnier et al. 2001; Wang et al. 2008; Wang et al. 2014) others assume elevations close to modern values by the latest at the middle Oligocene (Ding et al. 2014; Quade et al. 2011; Rowley & Currie 2006; Xu et al. 2013) or that a massive uplift occurred in the late Neogene (e.g., Molnar et al. 1993; Su et al. 2019; Wei et al. 2016).

During the last decade, a growing number of paleontological studies provide evidence for low elevated parts of Tibet until the early Neogene or even later. For example, the presence of subtropical to warm temperate floras during the late Eocene to early Miocene have been demonstrated for the basins of Hoh Xil, Kailas, Lunpola, Nima, and Qiabulin of southern and central parts of the Plateau (Ai et al. 2019; Ding et al. 2020; Miao et al. 2016; Su et al. 2019; Sun et al. 2014; Wu et al. 2017). These findings suggest that the present high-plateau character of Tibet with its dominant alpine environments is apparently a recent formation that did not emerge before the mid-Miocene. The young ages of species divergence in the phylogenies of high-altitude taxa endemic to the plateau are a logical consequence of – and evidence for – rather recent evolution of the TP (summary in Renner 2016). However, although it is becoming increasingly acknowledged that the HTO contributed to, and fostered, modern Asian biodiversity (Johansson et al. 2007; Steinbauer et al. 2016), our present concepts of the origin and historic biogeography of the terrestrial biotas inhabiting the HTO is far from being complete nor conclusive and has been hindered by a lack of and potential misinterpretation of data (Renner 2016; Spicer 2017; Spicer et al. 2020).

Phylogenies are a key mean in biogeographic and molecular evolutionary studies (Avise 2009; Avise et al. 2000) and increasingly recognized as being integral to research that aim to reconcile biological and geological information on landscape and biome in order to reconstruct Earth surface processes such as mountain building (Hoorn et al. 2013; Mulch & Chamberlain 2018). In fact, organismal evolution offers an independent line of evidence for the emplacement of major topographical features, which have been proved valid in refining the timing of events substantiated by geologic record. Specifically, several studies have demonstrated the suitability of phylogenetic data for addressing the timing and complexity of orogenic events, e.g., the Andean uplift and the formation of the Qinghai-Tibetan region (Antonelli et al. 2009; Luebert & Muller 2015).

We here use spiny frogs of the tribe Paini (Dicroglossidae) to untangling the spatiotemporal evolution of this group in the HTO and, thus, as an indirect indicator for the topographic and paleoecological development of High Asia. Spiny frogs are found across the Himalayan mountain arc from northern Afghanistan, Pakistan, and northern India, through Nepal, Sikkim, and Bhutan, and in the valleys of southern and eastern Tibet, eastwards to eastern China, and southwards to the mountains of Indochina (Myanmar, Thailand, Laos, northern Vietnam; Frost 2021). They live mostly in boulder-rich running water (Dubois 1975) or clear pools with flowing water. Males are characterized by black, keratinous spines. The Paini tribe is currently composed of the genus *Nanorana* Günther, 1896 (around 30 species), *Quasipaa* Dubois, 1992 (11 species), *Allopaa* Ohler and Dubois, 2006 (possibly two species), and the monotypic genus *Chrysopaa* Ohler and Dubois, 2006, with one species, *C. sternosignata* (Murray, 1885). Following Che et al. (Che et al. 2010) and our own findings (Hofmann et al. 2019), *Nanorana* can be subdivided into three subgenera (*Nanorana*, *Paa*, and *Chaparana*). However, the phylogenetic and mostly taxonomic relationships among Paini are not completely resolved with several taxonomic changes during the last decade including taxa descriptions (Che et al. 2009; Frost 2021; Huang et al. 2016; Jiang et al. 2005; Pyron & Wiens 2011).

Previous studies proposed contrasting hypotheses to explain the current distributional and phylogenetic patterns of spiny frogs in the HTO. While a strict vicariance driven scenario suggests species formation among major lineages when the species were “trapped” in the mountain mass and become separated when it uplifted (Che et al. 2010), a more recent study found no clear support for this model but indications for a Paleo-Tibetan origin of Himalayan spiny frogs (Hofmann et al. 2019), matching modern hypotheses for the past topographic surfaces of the southern parts of the HTO. This Tibetan-origin scenario (Schmidt et al. 2012) assumes that the ancestral lineages of Himalayan spiny frogs had adapted to the high-altitude environment in South Tibet, prior to the final uplift of the Greater Himalaya. With the continuously rising Himalayan mountain belt and the associated drying of southern Tibet, these ancestral lineages have probably been forced to track the displaced suitable environment along the transverse valleys of the Himalayas, such as the Brahmaputra, Kali Gandaki, or the Indus catchment. A hypothesis about the South-Tibetan origin has been also demonstrated in other Himalayan faunal elements, e.g., forest-dwelling *Pterostichus* ground beetles (Schmidt et al. 2012) and *Scutiger* lazy toads (Hofmann et al. 2017).

The phylogenetic placement of the most western Dicroglossid frogs that occur in the HTO (*Allopaa* from Kashmir Himalaya and *Chrysopaa* from Hindu Kush) has never been addressed. It is, however, of particular importance for a comprehensive understanding on how and when the Paini phylogeny has been shaped by the spatio-temporal surface uplift of the HTO. Therefore, we here reanalysed our previous dataset (Hofmann et al. 2019) by extending it with sequence data from *Allopaa* and *Chrysopaa.* We use our findings of the Paini phylogeny and time tree to discuss the biogeographic history of these frogs against the background of current HTO uplift concepts.

## Materials & Methods

### Sampling, laboratory protocols and data acquisition

We used sequence data of the 16S ribosomal, COI mitochondrial and Rag1 nuclear region available from our previous study (Hofmann et al. 2019) and complemented the data with a newly generated sequences for these three gene regions from *Allopaa hazarensis* (Dubois and Khan, 1979) (n = 6; Pakistan, including the type locality of the species – Datta, Manshera District, Hazera Division; for details see Fig. 1 and supplemental Tab. S1). Sampling was conducted according to the regulations for the protection of terrestrial wild animals under the permits of the Pakistan Museum of Natural History, Islamabad, Pakistan [No. PMNH/EST-1(89)/05]. We also included 16S rRNA and COI sequence data of *Chrysopaa sternosignata* from the Hindu Kush available in NCBI GenBank (accession numbers: MG700155 and MG699938). Our *Nanorana* samples from Himachal Pradesh, which were previously referred to as “sp.” (Hofmann et al. 2019), were identified as *Nanorana vicina* based on morphological characters (Boulenger 1920; Stoliczka 1872); for photos of live specimens Fig. S1). Genomic DNA was extracted from ethanol-muscle tissues using the DNeasy Blood & Tissue Kit (Qiagen, Venlo, Netherlands) following the manufacturer’s protocol. Approximately 571 bp of the ribosomal RNA (rRNA) 16S, 539 bp of the COI, and a sequence segment of 1207 bp of Rag1 gene were amplified via the polymerase chain reaction (PCR) using primers and PCR conditions as previously described (Hofmann et al. 2019). PCR products were purified using the mi-PCR Purification Kit (Metabion, Planegg, Germany) and the ExoSAP-IT enzymatic clean-up (USB Europe GmbH, Staufen, Germany; manufacturer’s protocol) or directly purified by Eurofins Genomics (Germany) with in-house protocols. The Sanger sequencing was performed on an ABI 3730 XL sequencer at Eurofins Genomics or by Macrogen Inc. (Seoul, South Korea or Amsterdam, Netherlands; http://www.macrogen.com).

**Figure 1:**
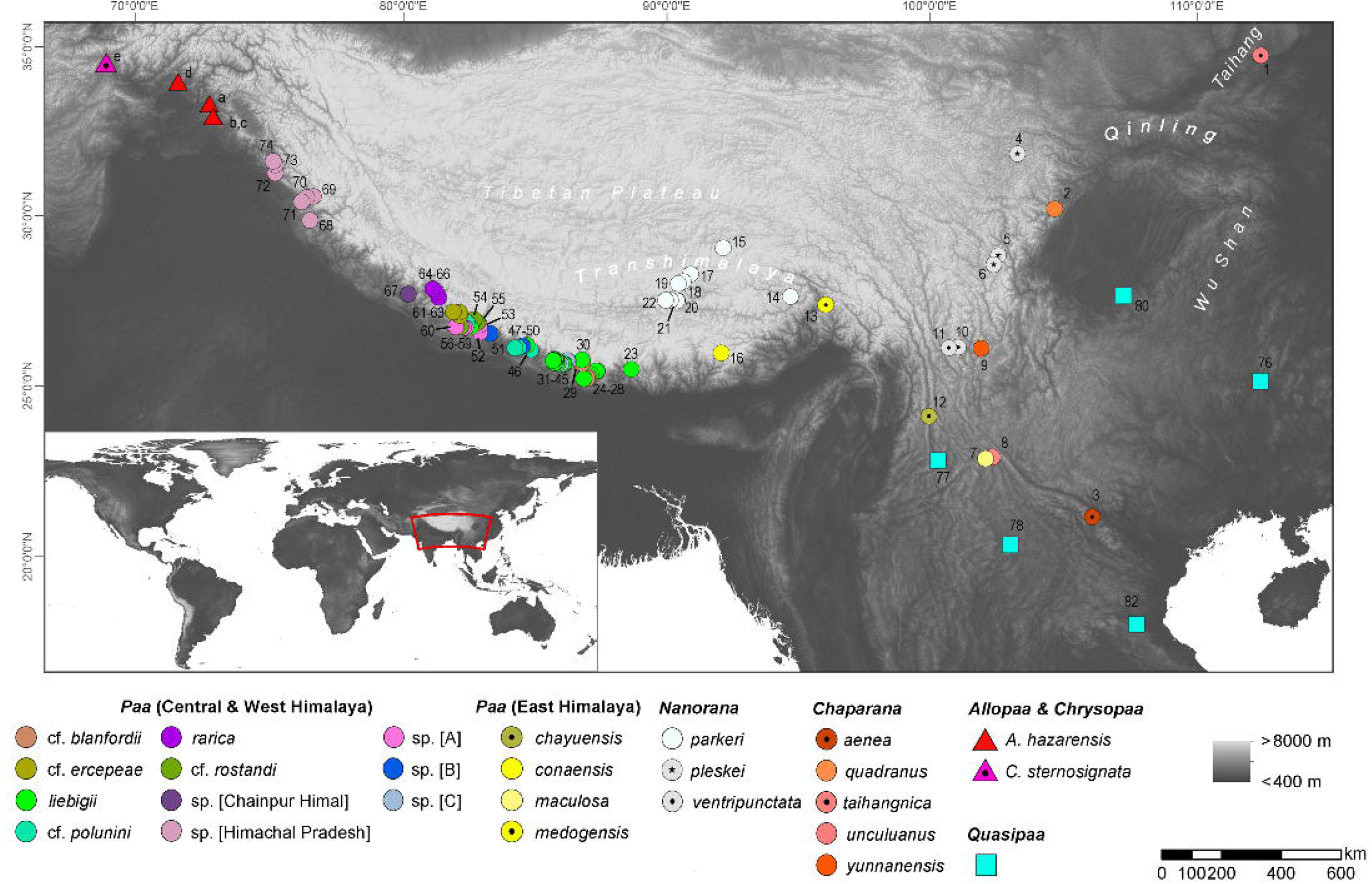
Map showing localities of sequences used in this study; locality numbers refer to samples/ sequences listed in supplemental Tab. S1.

### Sequence alignment and phylogenetic reconstruction

We aligned our new 16S sequences to the previous secondary structures-based data set (Hofmann et al. 2019) by eye; sequences of the protein-coding genes were aligned using the MUSCLE algorithm (Edgar 2004) in MEGA X (Kumar et al. 2018). Alignment based on nucleotides and amino acids produced similar results, since no ambiguities, such as deletions, insertions, or stop codons, were found.

The final concatenated rRNA + mtDNA + nuDNA sequence dataset consisted of 184 taxa and contained 2317 alignment positions of which 494 were phylogenetically informative. Nuclear data were unphased as most of the taxa had only single representative individuals. We inferred a maximum-likelihood (ML) and a Bayesian inference (BI) tree based on the concatenated sequence data using RAxML v.8.2.12 (Stamatakis 2014), IQ-TREE v.2.0 and MrBayes v.3.2.6 (Ronquist et al. 2012). The dataset was partitioned a priori by gene and codon fragments, and PartitionFinder 1.1.1 (Lanfear et al. 2012) was applied to optimize partitions using linked branch lengths, the corrected Aikaike Information Criterion (AICc), the greedy search algorithm, and the substitution models implemented in MrBayes and RAxML. We ran RAxML with the GTRGAMMA model and 1,000 bootstrap replicates on the CIPRES Cyberinfrastructure for Phylogenetic Research (Miller et al. 2010). IQ-TREE was performed with the edge-linked partition model (Chernomor et al. 2016) and both SH-like approximate likelihood ratio test (SH-aLRT) (Guindon et al. 2010) and the ultrafast bootstrap approximation (Hoang et al. 2018) using 1 Mio replicates per test. In the Bayesian analysis we assigned the doublet model (16×16) proposed by Schoniger and colleague (Schoniger & von Haeseler 1999) to the rRNA stem regions. For this procedure, unambiguous stem pairs were derived based on the consensus structure from RNAsalsa and specified in the MrBayes input file. For the analysis of the remaining positions, the standard 4×4 option was applied using a GTR evolutionary model for all nucleotide partitions. The site-specific rates were set variable. For reasons of comparison, we also inferred the Bayesian tree using the 4×4 standard model of DNA substitution for all regions and the optimized models and partitions as suggested by PartitionFinder. MrBayes was run for five million generations, sampling trees every 500th generation and using a random tree as a starting point. Inspection of the standard deviation of split frequencies after the final run as well as the effective sample size value of the traces using Tracer v. 1.7.1 (Rambaut et al. 2018) indicated convergence of Markov chains. In all analyses, four parallel Markov chain Monte Carlo simulations with four chains (one cold and three heated) were run. The first 25% of the samples of each run were discarded as burn-in. Based on the sampled trees, consensus trees were produced using the sumt command in MrBayes.

### Molecular dating

Divergence dates were estimated based on the full concatenated dataset, using BEAST2 v.2.6.2 (Bouckaert et al. 2014). Similar as to the MrBayes analyses, the partition scheme was optimized using PartitionFinder and the models that are implemented in BEAST. It is not possible to consider secondary structure information in BEAST (ambiguities are treated as unknown data so we did not remove stem regions) – thus all positions of the respective rRNA partition were treated under the same evolutionary model. Age constraints were derived from our previous calibration analysis of the phylogeny of *Nanorana,* which based on fossil-calibrated divergence estimates: MRCA of Paini 38.10 Ma, 28.70–47.50 (normal, sigma: 4.80); divergence of Tibetan *Nanorana* and Himalayan *Paa* 12.59 Ma, 7.93–17.30 (normal, sigma: 2.38; divergence Plateau frog *(Nanorana parkeri)* and *N. ventripunctata+N. pleskei* 6.35 Ma, 3.54–9.16 (normal, sigma: 1.44).

Analysis was based upon ten independent BEAST runs with a chain length of 50 million, a thinning interval of 5,000, a lognormal relaxed clock model, a Yule tree prior, a random tree as starting tree, and the site models selected using bModelTest package (Bouckaert & Drummond 2017) implemented in BEAST2. Runs were then combined with BEAST2 LogCombiner v.2.6.2 by resampling trees from the posterior distributions at a lower frequency, resulting in 9010 trees. Convergence and stationary levels were verified with Tracer. The final tree was obtained with TreeAnnotator v.2.6.2 and visualized with FigTree v. 1.4.3 (Drummond & Rambaut 2007).

## Results

### Phylogeny of Paini from the HTO

In both the ML and BI analyses, a relatively well resolved tree was obtained with strong support for most of the main clades, although with partly inconsistent and uncertain branching patterns of lineages within (sub)clades (Fig. 2). When information on secondary structure of 16S rRNA is considered (BI-tree), the results strongly support three monophyletic clades within Paini, apart from the monotypic *Chrysopaa*: *Quasipaa*, *Allopaa*, and *Nanorana*, with *Allopaa* forming the sister taxon to all *Nanorana*. Otherwise, *Allopaa* clusters with *Chaparana*, which together form the sister clade to *Paa* and *Nanorana* subgenera in the ML-tree (see also Fig. S2 for topology generated with IQ-TREE and with MrBayes using the 4×4 substitution model). The most striking result, consistently recovered in all trees, is the placement of *Chrysopaa* from the northern-central Afghanistan (southern slope of Hindu Kush), which forms the sister taxon to *Allopaa* and *Nanorana*.

**Figure 2:**
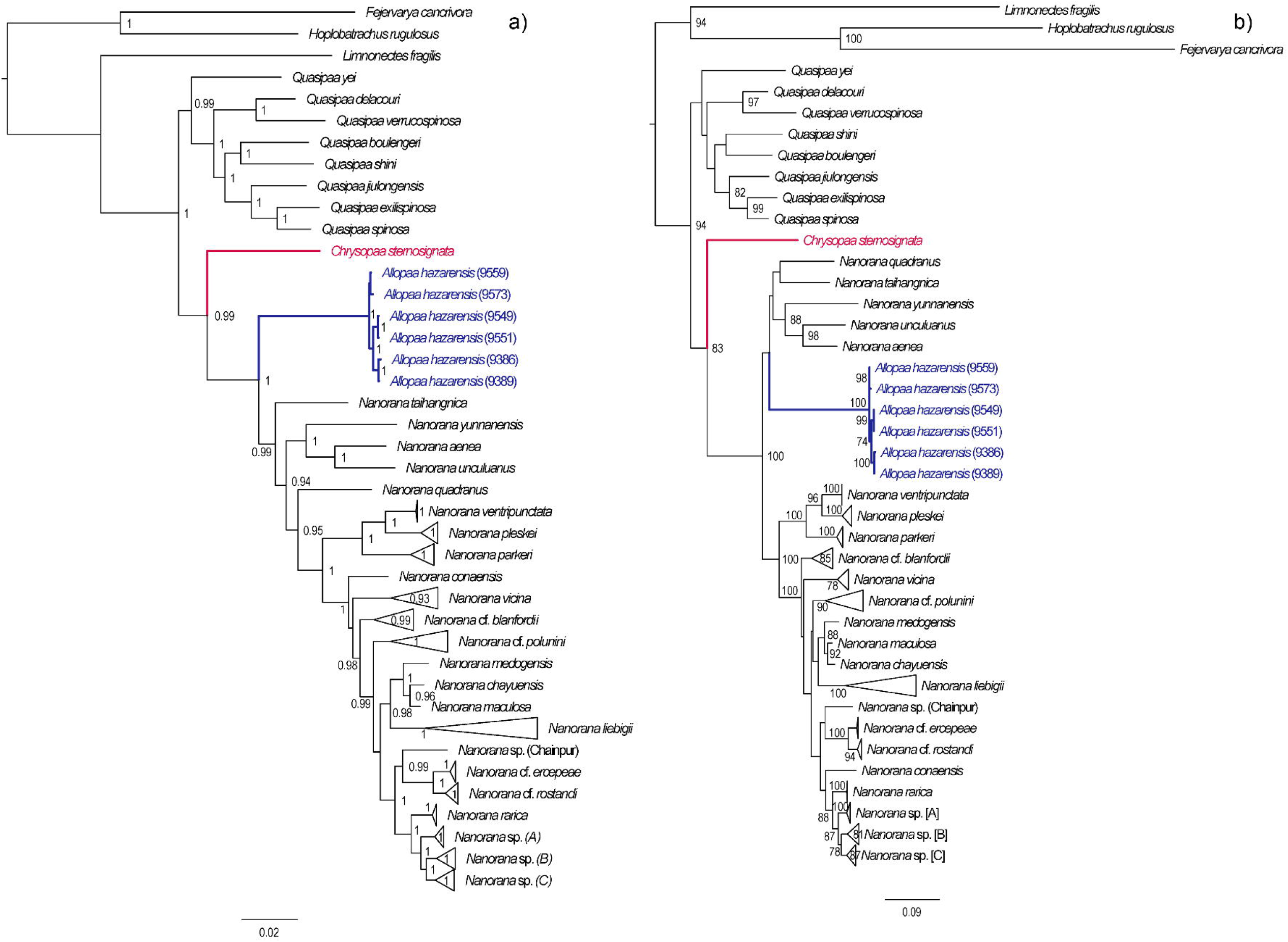
Bayesian inference (left) and Maximum-likelihood (right) tree based on concatenated mtDNA and nuDNA sequence data. Numbers at branch nodes refer to posterior probabilities ≥ 0.9 and bootstrap values > 70, respectively. For IQ-TREE topology see supplemental Fig. S2.

In accordance with our previous findings, three monophyletic subclades can be distinguished within *Nanorana,* namely *Chaparana* from montane regions of the southeastern margin of the TP and mountains of NE China, *Paa* from high-montane regions of the West, Central and East Himalaya, and nominal *Nanorana* from (sub)alpine regions of the TP and its eastern margin. Monophyly of *Chaparana* is not supported in the analyses if secondary structure of 16S is ignored. All *Paa* species together form the most species diverse clade.

Since the placement of *Allopaa* is of particular interest in terms of the origin and past biogeography of Paini, we tested the resulting topologies of major clades: BI tree considering secondary structure information of 16S, t_1_: *(Allopaa(Nanorana* genus)); RAxML/BI without secondary structure information, t2: ((*Chaparana*, *Allopaa*)(*Nanorana* sensu stricto, *Paa* sensu stricto)). We used a Bayes Factor (BF) approach and the tree topology tests implemented in IQ-TREE, namely the approximately unbiased (AU) test (Shimodaira 2002) as well as the RELL approximation (Kishino et al. 1990), including bootstrap proportion, Kishino-Hasegawa test (Kishino & Hasegawa 1989), Shimodaira-Hasegawa test (Shimodaira & Hasegawa 1999), and expected likelihood weights (Strimmer & Rambaut 2002). The marginal likelihoods estimations (MLE) for the BF calculations were obtained under each model based on both the stepping-stone (ss Xie et al. 2011)) and path sampling (ps Lartillot & Philippe 2006) methods implemented in BEAST v.1.10.4 (Suchard et al. 2018) using optimal partitions and substitution models as assessed in PartitionFinder, 250 million generations, logging interval of 25,000, a MLE chain length of 1 million, and 100 path steps. Statistical support was then evaluated via 2lnBF using the ps/ss results as per Kass & Raftery (Kass & Raftery 1995). Finally, we also used the stepping-stone approach with 10 million generations (4 runs and 4 chains), to estimate the model likelihood values for BF calculation with MrBayes by implementing the doublet option on 16S RNA stem regions and the standard substitution option on all other regions. We tested the hard constraint *vs.* negative constraint on *Chaparana* and *Allopaa*. In practice, any two models are compared to evaluate the strength of evidence against the null hypothesis (H0), defined as the one with the lower marginal likelihood (i.e., with the smaller value of the negative log-likelihood): 2lnBF < 2 indicates no evidence against H0; 2–6, weak evidence; 6–10, strong evidence; and > 10 very strong evidence. For the RELL approximation we used 1 Mio replicates, all other settings were left as default.

The AU test does not reject one of the two placement models for *Allopaa* (Tab. 1), as do the results of all other IQ-TREE tests. However, the BF of 28 (ss) and 32 (ps), based on the model likelihood values estimated with BEAST, strongly rejects a basal placement of *Allopaa* relative to the genus *Nanorana* in favor of the topology seen in the ML tree. Similarly, the marginal likelihoods calculated based on the runs considering the secondary structure of 16S were significantly higher for the unconstraint model (Tab. 1). Thus, the phylogenetic position of *Allopaa* as sister clade to *Chaparana* seems to be most likely, thereby making the *Nanorana* genus paraphyletic.

**Table 1:** Tree topology comparisons between the two models of *Allopaa* placements (t_1_, t_2_) based on Bayesian factor (BF) using BEAST, as well as the unbiased (AU) test (Shimodaira 2002), bootstrap proportion using RELL method (Kishino et al. 1990), Kishino-Hasegawa (KH) test (Kishino & Hasegawa 1989), Shimodaira-Hasegawa (SH) test (Shimodaira & Hasegawa 1999), and expected likelihood weights (ELW) using IQ-TREE; BF was also calculated for a hard constraint on *Chaparana* and *Allopaa* (A+Ch) *vs*. an unconstraint constellation using the stepping-stone approach in MrBayes and considering the secondary structure information of 16S. A = *Allopaa;* C = *Chaparana;* N = *Nanorana (genus);* P = *Paa;* ps = path sampling log marginal likelihood; ss = stepping-stone log marginal likelihood; + = a tree is not rejected if its p-value > 0.05. Bold log marginal likelihood values indicate the model most favored by a method (higher is better).

| Topology | ps | ss | 2lnBF | logL | deltaL | bp-RELL | p-KH | p-SH | c-ELW | p-AU |
| --- | --- | --- | --- | --- | --- | --- | --- | --- | --- | --- |
| t<sub>1</sub>(A(N)) | -15471 | -15477 | ps: 32 | -14164.109 | 2.458 | 0.397+ | 0.383+ | 0.383+ | 0.399+ | 0.383+ |
| t<sub>2</sub>((A+Ch)(P,N)) | **-15455** | **-15463** | ss: 28 | **-14161.652** | 0 | 0.603+ | 0.617+ | 1+ | 0.601+ | 0.617+ |
| (A+Ch) |  | **-16472** | ss: 56 |  |  |  |  |  |  |  |
| unconstraint |  | -16500 |  |  |  |  |  |  |  |  |

### Divergence times in spiny frogs

Dating analysis suggests an origin of Paini *(Allopaa, Chrysopaa, Nanorana, Quasipaa)* in the mid Oligocene (28.21 Ma, 20.11-35.18 Ma), what is in the range of previous estimations (Che et al. 2010; Hofmann et al. 2019; Sun et al. 2018) (Fig. 3). The age of Himalayan-Tibetan spiny frogs *(Allopaa, Chrysopaa, Nanorana)* is estimated to be 25.7 Ma (18.70-32.16). Interestingly, within crown *Allopaa*+*Nanorana*, the clade comprising the montane *Chaparana* and West-Himalayan *Allopaa* split from the Central/East Himalayan and Tibetan *Nanorana* (subgenera *Paa* and *Nanorana*) in the early Miocene, around 20 Ma, followed by the separation of *Chaparana* and *Allopaa ca.* 3 million years later. The divergence of the nominal *Nanorana* (endemic to the TP) from *Paa* (Greater Himalaya) occurred around 15 Ma (11.45– 18.27 Ma). This estimate is close to the age of 13 Ma (7–25 Ma) calculated by Sun et al. (Sun et al. 2018), and 10–12 Ma estimated by Wiens et al. (Wiens et al. 2009).

**Figure 3:**
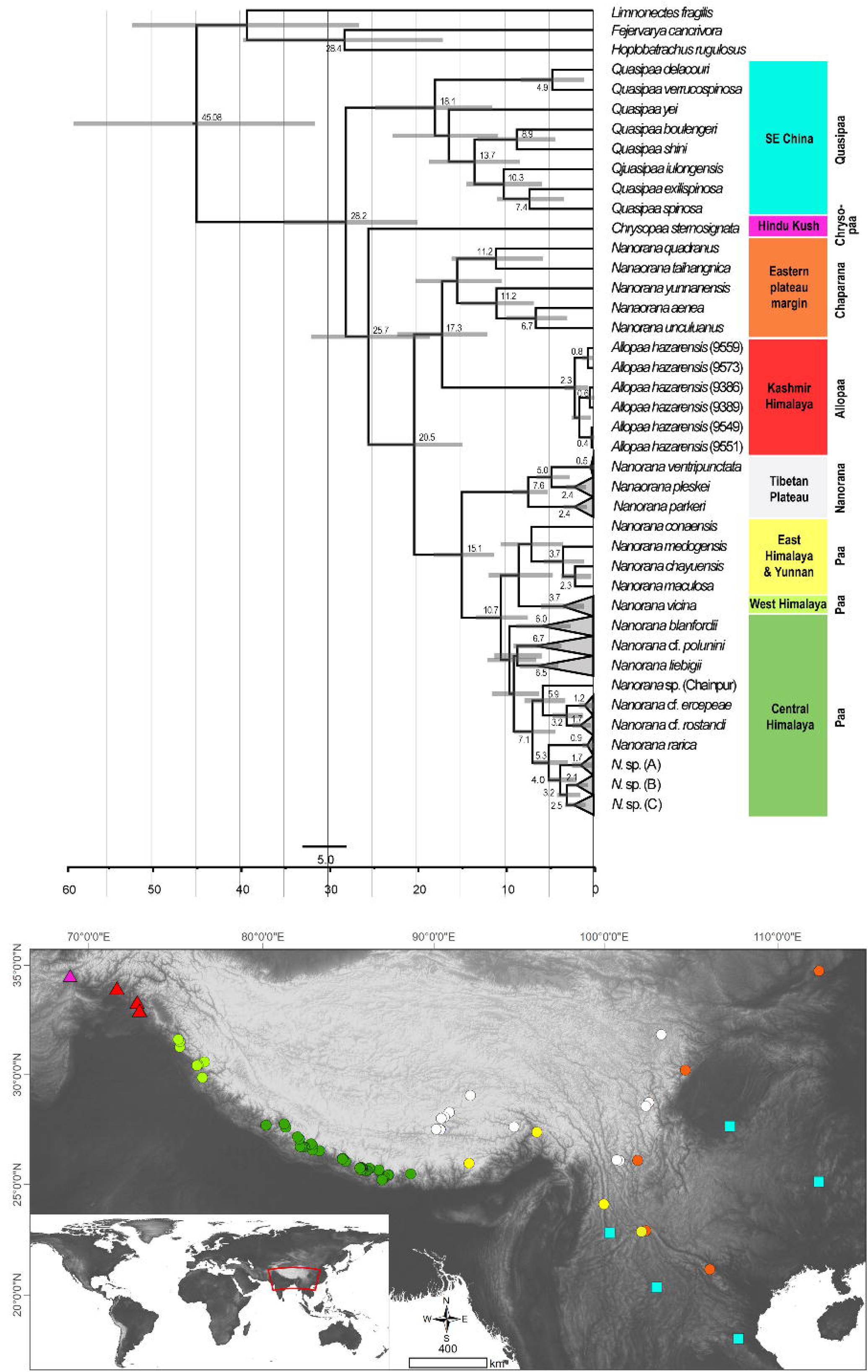
Ultametric time-calibrated phylogenetic tree obtained with BEAST2 for the concatenated sequence data in spiny frogs. Grey bars indicated the 95% HPD for the respective nodes; ages are shown for nodes that are supported by Bayesian posterior probability ≥ 0.95.

Diversification of Central Himalayan *Paa* clades has taken place continuously during the whole Mid to Late Miocene. Most of the main lineages within *Paa* were present at least in the late Miocene, and nearly all species are not younger than the Pliocene.

## Discussion

We here report the first, well-supported phylogeny of the westernmost HTO Paini taxa *Chrysopaa sternosignata* and *Allopaa hazarensis* in the context of their closest relatives. Our work based on sequence information of *A. hazarensis* specimens from the foothills of the Kashmir Himalaya, a previously published data set (Hofmann et al. 2019), and additional sequence data of *C. sternosignata* from the southern foothills of the Hindu Kush in Afghanistan available from GenBank. The study provides evidence for an early-Miocene evolution of Himalayan Paini, which is ultimately linked to the paleoecological evolution of the HTO.

Consistent with our previous results (Hofmann et al. 2019), the Southeast Asian genus *Quasipaa* is sister to all other spiny frogs. Most remarkable, the monotypic *Chrysopaa* is placed basally relative to *Nanorana* and *Allopaa,* supporting the presence of ancestral Paini lineages in the far north western part of the HTO, which is diametrically opposite end of the HTO with respect to the ancestral area of spiny frogs that is assumed to be the Paleogene East or Southeast China (Che et al. 2010; Hofmann et al. 2019). Thus, it can be assumed that the ancestor of *Chrysopaa* appeared elsewhere near the eastern margin of the HTO during the late Oligocene-early Miocene. If so, it implies that members of the *Chrysopaa* stem group must have been temporarily present in the interior of the HTO during the following time, to enable a range expansion up to the western margin of the mountains system. Given this scenario, the climatic preferences of ancestral spiny frogs are of particular interest. Most anurans show remarkable stasis in ecological niches, suggesting that dispersal will have been historically constrained between similar climatic conditions (Wiens 2011). Since all species of the most basal clade *Quasipaa* are adapted to the subtropical climate, a similar temperature preference must be assumed for the *Chrysopaa* ancestor. This preference has not changed significantly during the Neogene period as *C. sternosignata* occurs under subtropical to warm temperate climate conditions in the colline zone south of the Hindu Kush (Pakistani Balochistan) and the Kashmir valley (Khan 2006; Sarwar et al. 2016; Wagner et al. 2016). Consequently, a subtropical climate associated with sufficient humidity suitable for amphibians must have existed in large parts of the late Oligocene-Tibet to allow a trans-Tibetan dispersal of *Chrysopaa* stem group members. Interestingly, basal divergences of West Himalayan taxa are also known from the gekkonid genus *Cyrtodactylus*, dating even back to the early Eocene (Agarwal et al. 2014).

Also unexpected are our results with respect to the phylogenetic position and timing of the evolution of *Allopaa* from the foothills of the Kashmir Himalaya. This group evolved during the early to mid-Miocene most parsimoniously as sister clade to *Chaparana.* Species of the latter taxon occur along the eastern margin of the HTO and therewith at the opposite end of the HTO where *Allopaa* is distributed. *Chaparana* and *Allopaa* together constitute the sister clade to the Tibetan *Nanorana* and Himalayan *Paa*, which indicates on the one hand that *Nanorana* might be paraphyletic with respect to *Allopaa.* On the other hand, it shows that *Allopaa* is phylogenetically not related to the biogeographically neighboring Himalayan spiny frogs. This finding is crucial with respect to the ancestral distributional area of the *Chaparana+Allopaa* clade and their ancestral habitat preferences. Recent species of *Chaparana* occur in the colline and lower montane zone along the eastern margin of the HTO and the easterly neighbored mountains and, thus, immediately adjacent to (or overlapping with) the supposed ancestral area of spiny frogs (Che et al. 2010; Hofmann et al. 2019). Similar as assumed for *Chrysopaa,* the ancestor of *Allopaa* must have been dispersed across a moderately elevated Tibetan Plateau, although about eight million years later than the ancestor of *Chrysopaa.* Since species of *Allopaa* occur under warm-temperate conditions in the colline to lower montane zone (comparable to those of its sister group *Chaparana)* (Ahmed et al. 2020), similar temperature preferences can be assumed for ancestral *Allopaa*. Therefore, the supposed trans-Tibet dispersal event of this lineage implies the presence of warm temperate conditions in significant parts of Tibet’s interior at least up to the early-mid Miocene boundary. Due to the progressive uplift of Tibet and the associated continuous cooling, the *Allopaa* stem group members might have successively been lost to extinction. Today’s absence of members of *Chaparana* and *Allopaa* in the high montane zone throughout the HTO suggests that species of their ancestral lineages were not able to adapt fast enough to the new conditions under a dramatically changing environment. Alternatively, a westward and northwestward spread of ancestral *Allopaa* along the southern slopes of the Himalaya must also be considered. However, this model is very unlikely, as it would imply extinction of all ancestral lineages in fast areas covering almost the whole Himalayan mountain arc.

Considering that since the onset of surface uplift subtropical to warm temperate environments have always been present along the Himalayan southern slope, such radical extinction or turnover is implausible given the recent and former ecological conditions in this area. Moreover, the absence of *Allopaa* species, but occurrence of many spiny frogs of the subgenus *Paa* along the southern slopes of the eastern, central, and western Himalaya and north to the Indian Himachal Pradesh, contradicts this extinction scenario.

Unlike spiny frogs of the taxa *Chrysopaa, Allopaa* and *Chaparana* which are restricted to the subtropical to warm temperate climate, many representatives of the monophyletic *Nanorana*+*Paa* clade are adapted to climatically colder habitats and known to occur in the high montane and subalpine-alpine zones of the HTO. The evolutionary late appearance of this clade is indicative for the minimum age of high-altitude environments in the HTO: Although spiny frogs were present in the area since at least the early Paleogene/Neogene boundary, cold-adapted species did not evolve before *ca.* 15 Ma (Fig. 3). This is a strong hint that extensive high-altitude environments were present in the HTO only from mid-Miocene at earliest.

## Conclusions

We provide the first phylogeny of the most western Himalayan spiny frogs. Our findings suggest a late Oligocene to early Miocene dispersal of two subtropical/temperate lineages, *Chrysopaa* and *Allopaa,* from the ancestral area of spiny frogs in SE Asia across the HTO into its far north western part. This dispersal scenario is central to the long-standing debate regarding the paleoenvironmental and paleoelevational development of the TP. Given the stem age of subtropical *Chrysopaa* of ca. 26 Mya and the divergence time of 17 Mya between warm temperate *Allopaa* and *Chaparana*, the results strongly indicate the large-scale presence of subtropical environments north of the present Himalayas until the late Oligocene, and of warm temperate climates until the late Miocene. This contrasts with known geoscientific models of the paleoeleventional evolution of the TP which assume large scale surface uplift close to present heights until the mid-Oligocene (e.g., Kapp et al. 2007; Mulch & Chamberlain 2006; Tapponnier et al. 2001; Wang et al. 2008; Wang et al. 2014). However, over the last decade a growing number of fossil data provide evidence for the presence of tropical to warm temperate floras and freshwater fishes in central Tibet during the late Paleogene until the early Neogene (Song et al. 2010; Su et al. 2019; Wei et al. 2016; Wu et al. 2017). Consistent with these findings our results support the recent concept proposed by Spicer and colleagues (Spicer et al. 2020), which assumes that the TP was not uplifted as a whole, but instead, a deep wide east–west oriented valley occurred in the Tibetan interior before the final plateau formation. We suspect that this supposed valley represents the migration corridor of the ancestral *Chrysopaa* and *Allopaa* lineages, which today are represented by the two relict taxa, *C. sternosignata* and *A. hazarensis*, endemic to the region south of the Hindu Kush and Kashmir Himalaya. This scenario is in line with and adds to the Tibetan-origin hypothesis of the paleo-Tibetan fauna (Hofmann et al. 2017; Hofmann et al. 2019; Schmidt et al. 2013). Disjunct distribution patterns of species groups between the eastern and western part of the HTO, as we demonstrate here for spiny frogs, have been also observed in Broscini ground beetles, with the genus *Eobroscus* widely distributed in East Asia and Indochina and with *Kashmirobroscus* endemic to a small part of the Kashmir Himalaya (Schmidt et al. 2013). Moreover, the Kashmir Himalaya is well-known for the occurrence of several highly endemic ground beetles (Schmidt et al. 2012). We expect that numerous additional lineages endemic to the Kashmir Himalaya will be identified in future which may contribute to resolve the evolution of the HTO. We therefore encourage further and systematic research in this area and the use of more powerful molecular data, for example, through the use of genomic sequencing to better understand the evolution and Cenozoic history of Himalayan biodiversity against the background of existing geological scenarios.

## Acknowledgements

We thank Sandra Kukowa, Anja Bodenheim, and Jana Poláková for technical support in the lab. This work was funded by the German Research Foundation (DFG, grant no. HO 3792/8-1 to SH), and by the Slovak Research and Development Agency (grant no. APVV-19-0076 to DJ).

