## Supplemental Figure S1 for "Relict groups of spiny frogs indicate Late Paleogene-Early Neogene trans-Tibet dispersal of thermophile faunal elements"

**Supplemental Information Figure S1**

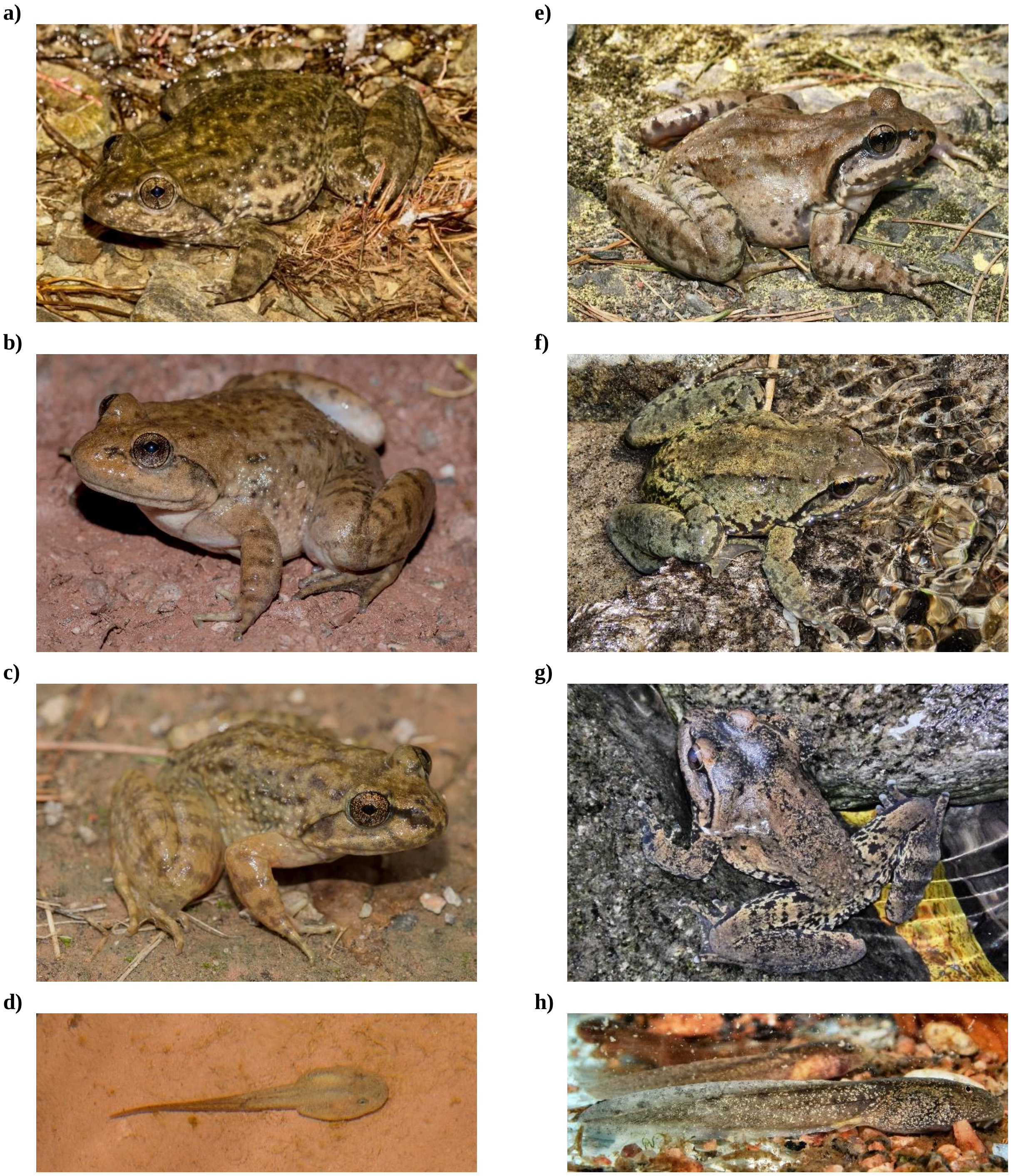


**Photo vouchers of *Allopaa hazarensis* (left panels) and *Nanorana vicina* (right panels)** from the present study, sampled for DNA in Pakistan and Himachal Pradesh, respectively (Photographs of *Allopaa hazarensis*: D. Jablonski; of *Nanorana vicina*: S. Litvinchuk): **a)** *A. hazarensis* from the type locality Datta, Pakistan (1300 m; locality no. a); **b)** *A. hazarensis* from Margi, Murree, Pakistan (1618 m; locality no. c); **c)** *A. hazarensis* from locality Laram Qilla, Lower Dir, Pakistan (1436 m; locality no. d); d) tadpole of *A. hazarensis* from Margi, Murre, Pakistan (1618 m, locality no. c); **e)** *N. vicina* from Narkanda (2650 m; locality no. 68); **f)** *N. vicina* from Pulga, (2199 m; locality no. 69); **g)** *N. vicina* from Panjpula (2016 m; locality no. 72); **h)** tadpole of *N. vicina* from Panjpula (2016 m; locality no. 72).
