## Supplemental Figure S2 for "Relict groups of spiny frogs indicate Late Paleogene-Early Neogene trans-Tibet dispersal of thermophile faunal elements"

**Supplemental Information Fig. S2**

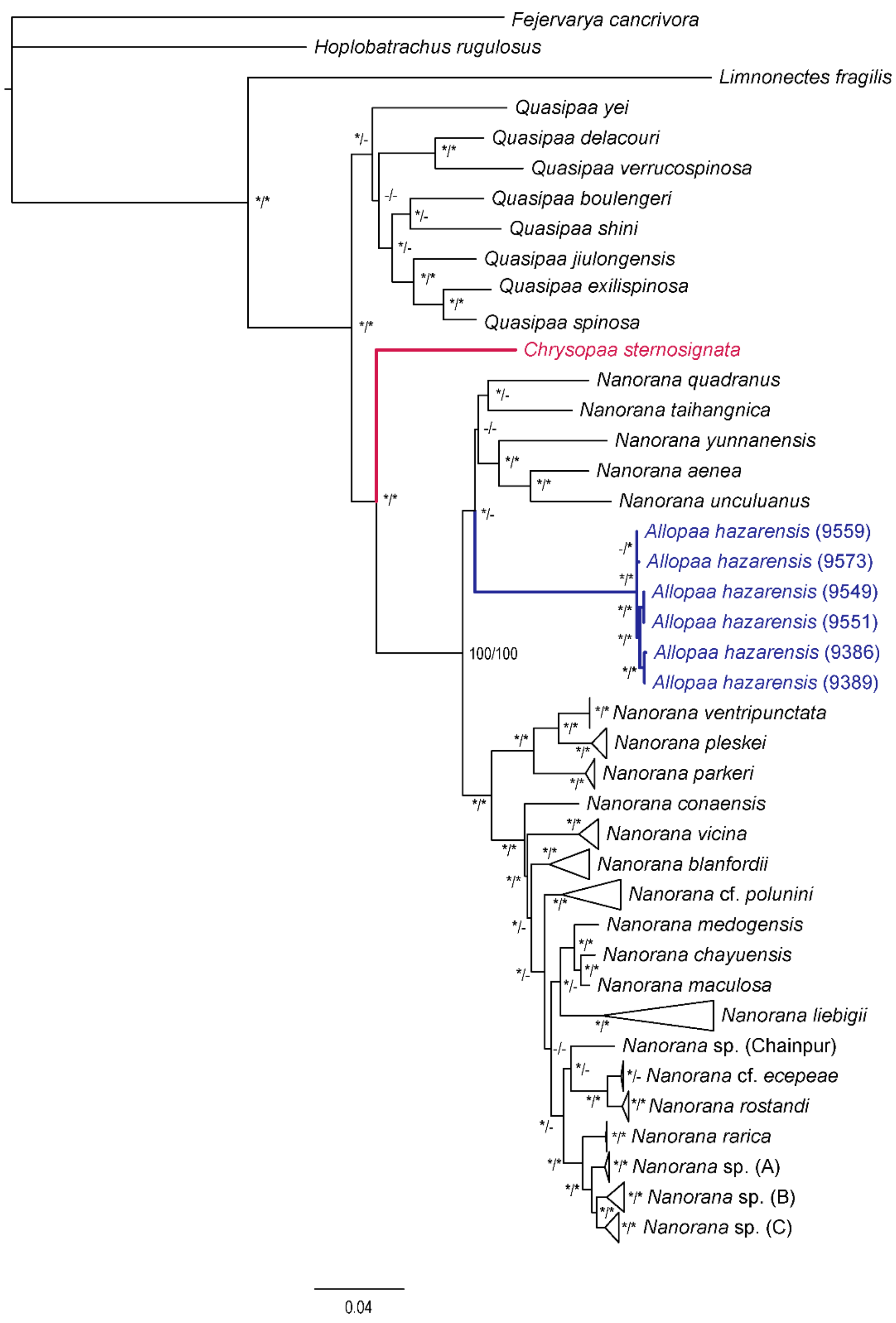


**IQ-TREE topology.** SH-aLRT (*≥70) and ultrafast bootstrap support (*≥ 90) values are indicated with an asterisk at the respective node.

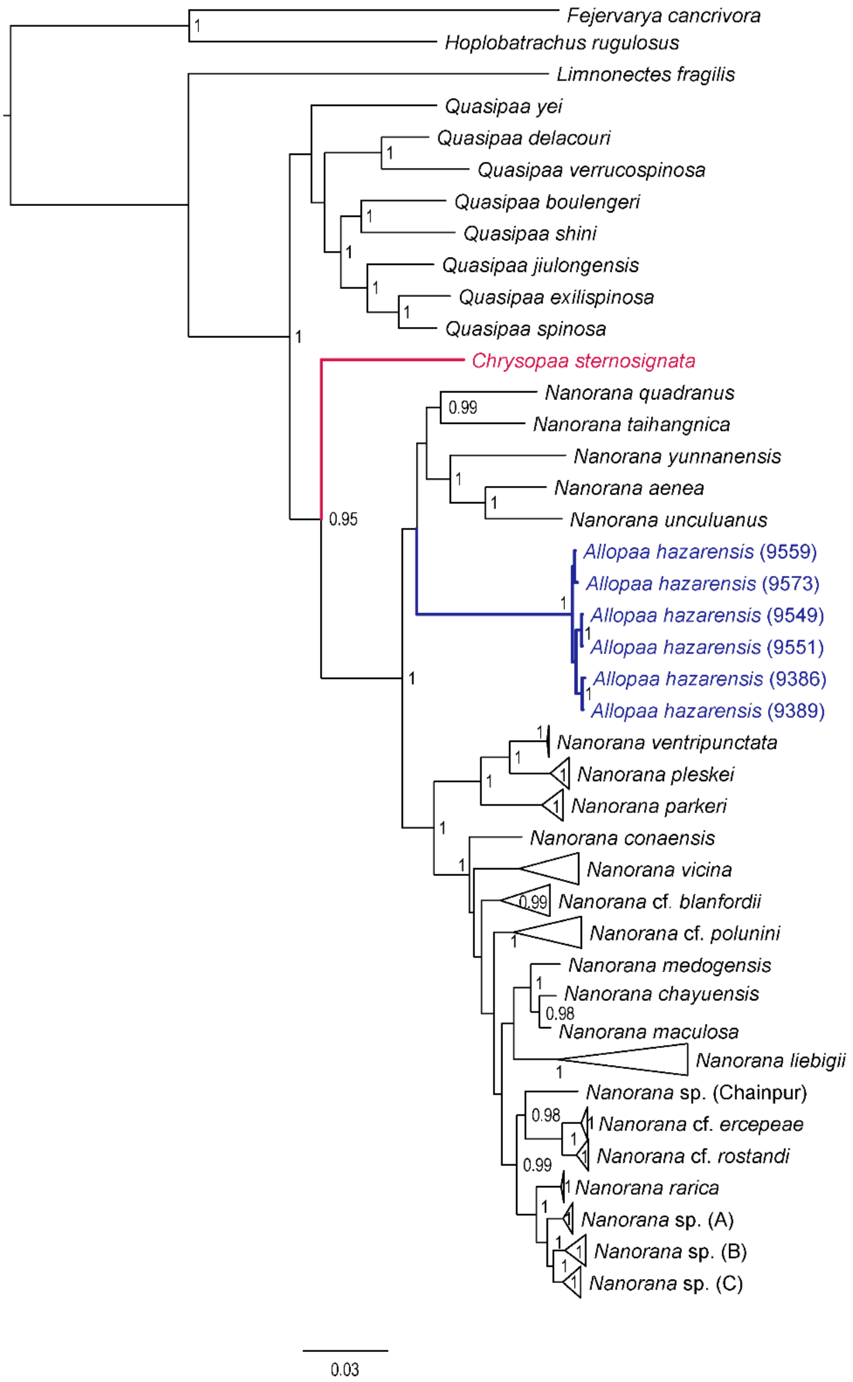


**MrBayes tree generated with the standard 4×4 model of DNA substitution.** Posterior probabilities ≥ 0.90 are shown at the respective node.
