## Supplemental Table S1 for "Relict groups of spiny frogs indicate Late Paleogene-Early Neogene trans-Tibet dispersal of thermophile faunal elements"

**Supplemental Information Tab. S1**

**List of species used in the present study, including sample ID or voucher numbers, sample localities and GenBank accession numbers.** Locality identifier (Loc) refer to the Fig. 1. Coordinates are given in decimal degrees. § = samples of photographed specimens (see Fig. S1); * = sample localities not shown in Fig. 1 because they lie far to SO China; for reference see map in Hofmann et al. (2019).

| Taxon | SampleID/voucher | Data origin | Loc | N | E | 16S | co1 | rag1 |
|---|---|---|---|---|---|---|---|---|
| *Allopaa hazarensis* | 9386 | This study | a | 34.295 | 73.258 | MW598397 | MW603002 | MW598465 |
| *Allopaa hazarensis* | 9389 | This study | a | 34.295 | 73.258 | MW598398 | MW603003 | MW598466 |
| *Allopaa hazarensis* | 9549 | This study | b | 33.951 | 73.468 | MW598393 | MW603004 | MW598461 |
| *Allopaa hazarensis* | 9551 | This study | c | 33.940 | 73.465 | MW598394 | MW603005 | MW598462 |
| *Allopaa hazarensis* | 9559 | This study | d | 34.785 | 71.986 | MW598395 | MW603006 | MW598463 |
| *Allopaa hazarensis* | 9573 | This study | d | 34.785 | 71.986 | MW598396 | MW603007 | MW598464 |
| *Chrysopaa sternosignata* | USNM:Herp589844 | NCBI | e | 34.940 | 69.255 | MG700155 | MG699938 | — |
| *N. aenea* | | NCBI | 3 | 22.336 | 103.844 | EU979830 | KR087830 | HM163609 |
| *N.* cf. *blanfordii* | JS040529_NME | Hofmann et al. 2019 | 29 | 27.617 | 87.233 | MN012067 | — | MN032491 |
| *N.* cf. *blanfordii* | JS040531_NME | Hofmann et al. 2019 | 29 | 27.617 | 87.233 | MN012068 | — | MN032492 |
| *N.* cf. *blanfordii* | JS040532_NME | Hofmann et al. 2019 | 29 | 27.617 | 87.233 | MN012069 | — | MN032493 |
| *N.* cf. *blanfordii* | JS040533_NME | Hofmann et al. 2019 | 29 | 27.617 | 87.233 | MN012070 | — | MN032494 |
| *N.* cf. *blanfordii* | JS040534_NME | Hofmann et al. 2019 | 29 | 27.617 | 87.233 | MN012071 | | MN032495 |
| *N.* cf. *blanfordii* | JS040535_NME | Hofmann et al. 2019 | 29 | 27.617 | 87.233 | MN012072 | — | MN032496 |
| *N.* cf. *blanfordii* | JS060520_NME | Hofmann et al. 2019 | 28 | 27.173 | 87.421 | MN012073 | — | MN032497 |
| *N.* cf. *blanfordii* | JS060515_NME | Hofmann et al. 2019 | 27 | 27.214 | 87.463 | MN012074 | — | MN032498 |
| *N.* cf. *blanfordii* | JS060508_NME | Hofmann et al. 2019 | 25 | 27.413 | 87.734 | MN012075 | — | MN032499 |
| *N. cf. ercepeae* | A2016/13_NME | Hofmann et al. 2019 | 67 | 29.374 | 81.137 | — | MN012211 | — |
| *N. cf. ercepeae* | A2017/13_NME | Hofmann et al. 2019 | 67 | 29.374 | 81.137 | MN012076 | MN012212 | MN032500 |
| *N. cf. ercepeae* | A1_12_NME | Hofmann et al. 2019 | 63 | 28.963 | 82.857 | MN012077 | MN012213 | MN032501 |
| *N. cf. ercepeae* | A7_12_NME | Hofmann et al. 2019 | 62 | 28.855 | 82.961 | MN012078 | MN012214 | MN032502 |
| *N. cf. ercepeae* | A4_12_NME | Hofmann et al. 2019 | 61 | 28.857 | 82.976 | MN012079 | MN012215 | MN032503 |
| *N. cf. ercepeae* | A5_12_NME | Hofmann et al. 2019 | 61 | 28.857 | 82.976 | MN012080 | MN012216 | MN032504 |
| *N. cf. ercepeae* | A6_12_NME | Hofmann et al. 2019 | 61 | 28.857 | 82.976 | MN012081 | MN012217 | MN032505 |
| *N.* cf. *polunini* | R15_12_NME | Hofmann et al. 2019 | 59 | 28.502 | 83.129 | MN012082 | MN012218 | MN032506 |
| *N.* cf. *polunini* | R20_12_NME | Hofmann et al. 2019 | 57 | 28.513 | 83.255 | MN012083 | — | MN032507 |
| *N.* cf. *polunini* | SH070507_NME | Hofmann et al. 2019 | 50 | 28.060 | 85.294 | MN012084 | MN012219 | MN032508 |
| *N.* cf. *polunini* | SH070509_NME | Hofmann et al. 2019 | 49 | 28.080 | 85.295 | MN012085 | MN012220 | MN032509 |
| *N.* cf. *polunini* | SH070531_NME | Hofmann et al. 2019 | 46 | 27.965 | 85.472 | MN012086 | MN012221 | MN032510 |
| *N.* cf. *polunini* | R3_09_13_NME | Hofmann et al. 2019 | 51 | 28.380 | 84.065 | MN012087 | MN012222 | MN032511 |
| *N.* cf. *rarica* | A1961/13_NME | Hofmann et al. 2019 | 66 | 29.510 | 82.090 | MN012202 | MN012322 | — |
| *N.* cf. *rarica* | A1970/13_NME | Hofmann et al. 2019 | 66 | 29.510 | 82.090 | MN012203 | MN012323 | MN032606 |
| *N.* cf. *rarica* | A2015/13_NME | Hofmann et al. 2019 | 66 | 29.510 | 82.090 | MN012204 | MN012324 | MN032607 |
| *N.* cf. *rarica* | A2019/13_NME | Hofmann et al. 2019 | 66 | 29.510 | 82.090 | MN012205 | MN012325 | MN032608 |
| *N.* cf. *rarica* | A1965/13_NME | Hofmann et al. 2019 | 65 | 29.513 | 82.092 | MN012206 | MN012326 | MN032609 |
| *N.* cf. *rarica* | A1960/13_NME | Hofmann et al. 2019 | 64 | 29.360 | 82.200 | MN012207 | MN012327 | — |
| *N.* cf. *rostandi* | R1_12_NME | Hofmann et al. 2019 | 60 | 28.513 | 83.033 | MN012088 | MN012223 | MN032512 |
| *N.* cf. *rostandi* | R2_12_NME | Hofmann et al. 2019 | 60 | 28.513 | 83.033 | MN012089 | MN012224 | MN032513 |
| *N.* cf. *rostandi* | R3_12_NME | Hofmann et al. 2019 | 60 | 28.513 | 83.033 | MN012090 | MN012225 | MN032514 |
| *N.* cf. *rostandi* | R4_12_NME | Hofmann et al. 2019 | 60 | 28.513 | 83.033 | MN012091 | MN012226 | MN032515 |
| *N.* cf. *rostandi* | R11_12_NME | Hofmann et al. 2019 | 59 | 28.502 | 83.129 | MN012092 | MN012227 | MN032516 |

| | | | | | | | | |
|---|---|---|---|---|---|---|---|---|
| *N.* cf. *rostandi* | R12_12_NME | Hofmann et al. 2019 | 59 | 28.502 | 83.129 | MN012093 | MN012228 | MN032517 |
| *N.* cf. *rostandi* | R13_12_NME | Hofmann et al. 2019 | 59 | 28.502 | 83.129 | MN012094 | MN012229 | MN032518 |
| *N.* cf. *rostandi* | R14_12_NME | Hofmann et al. 2019 | 59 | 28.502 | 83.129 | MN012095 | MN012230 | MN032519 |
| *N.* cf. *rostandi* | R16_12_NME | Hofmann et al. 2019 | 59 | 28.502 | 83.129 | MN012096 | MN012231 | MN032520 |
| *N.* cf. *rostandi* | R6_12_NME | Hofmann et al. 2019 | 59 | 28.502 | 83.129 | MN012097 | MN012232 | MN032521 |
| *N.* cf. *rostandi* | R7_12_NME | Hofmann et al. 2019 | 59 | 28.502 | 83.129 | MN012098 | MN012233 | MN032522 |
| *N.* cf. *rostandi* | R8_12_NME | Hofmann et al. 2019 | 59 | 28.502 | 83.129 | MN012099 | MN012234 | MN032523 |
| *N.* cf. *rostandi* | R9_12_NME | Hofmann et al. 2019 | 59 | 28.502 | 83.129 | MN012100 | MN012235 | MN032524 |
| *N.* cf. *rostandi* | R17_12_NME | Hofmann et al. 2019 | 56 | 28.519 | 83.264 | MN012101 | MN012236 | MN032525 |
| *N.* cf. *rostandi* | SH070550_NME | Hofmann et al. 2019 | 55 | 28.683 | 83.591 | MN012102 | — | MN032526 |
| *N.* cf. *rostandi* | SH070538_NME | Hofmann et al. 2019 | 54 | 28.680 | 83.594 | MN012103 | — | MN032527 |
| *N. chayuensis* | SCUM050410CHX | NCBI | 12 | 25.823 | 98.858 | EU979838 | — | HM163587 |
| *N. conaensis* | KIZ-YP152 | NCBI | 16 | 27.991 | 91.957 | EU979834 | — | HM163589 |
| *N. liebigii* | A17_12_NME | Hofmann et al. 2019 | 56 | 28.519 | 83.264 | MN012104 | MN012237 | MN032528 |
| *N. liebigii* | R18_12_NME | Hofmann et al. 2019 | 56 | 28.519 | 83.264 | MN012105 | MN012238 | MN032529 |
| *N. liebigii* | SH070515_NME | Hofmann et al. 2019 | 47 | 28.099 | 85.317 | MN012106 | — | MN032530 |
| *N. liebigii* | SH0805109_NME | Hofmann et al. 2019 | 45 | 27.673 | 86.240 | MN012107 | — | MN032531 |
| *N. liebigii* | SH080506_NME | Hofmann et al. 2019 | 42 | 27.609 | 86.295 | MN012108 | — | MN032532 |
| *N. liebigii* | SH080554_NME | Hofmann et al. 2019 | 41 | 27.718 | 86.311 | MN012109 | — | MN032533 |
| *N. liebigii* | SH080536_NME | Hofmann et al. 2019 | 38 | 27.691 | 86.343 | MN012110 | MN012239 | — |
| *N. liebigii* | SH080537_NME | Hofmann et al. 2019 | 38 | 27.691 | 86.343 | MN012111 | MN012240 | MN032534 |
| *N. liebigii* | SH080538_NME | Hofmann et al. 2019 | 38 | 27.691 | 86.343 | MN012112 | — | MN032535 |
| *N. liebigii* | SH080524_NME | Hofmann et al. 2019 | 37 | 27.694 | 86.351 | MN012113 | MN012241 | MN032536 |
| *N. liebigii* | SH080534_NME | Hofmann et al. 2019 | 37 | 27.694 | 86.351 | MN012114 | — | MN032537 |
| *N. liebigii* | Ne16_13_NME | Hofmann et al. 2019 | 36 | 27.584 | 86.411 | MN012115 | MN012242 | MN032538 |
| *N. liebigii* | Ne17_13_NME | Hofmann et al. 2019 | 36 | 27.584 | 86.411 | MN012116 | MN012243 | MN032539 |
| *N. liebigii* | Ne12_13_NME | Hofmann et al. 2019 | 34 | 27.584 | 86.594 | MN012117 | MN012244 | MN032540 |
| *N. liebigii* | Ne10_13_NME | Hofmann et al. 2019 | 33 | 27.586 | 86.635 | MN012118 | MN012245 | MN032541 |
| *N. liebigii* | JS040512_NME | Hofmann et al. 2019 | 30 | 27.631 | 87.224 | MN012119 | MN012246 | MN032542 |
| *N. liebigii* | JS040513_NME | Hofmann et al. 2019 | 30 | 27.631 | 87.224 | MN012120 | MN012247 | MN032543 |
| *N. liebigii* | JS060518_NME | Hofmann et al. 2019 | 28 | 27.173 | 87.421 | MN012121 | — | MN032544 |
| *N. liebigii* | JS060511_NME | Hofmann et al. 2019 | 26 | 27.296 | 87.535 | MN012122 | — | MN032545 |
| *N. liebigii* | JS060509_NME | Hofmann et al. 2019 | 25 | 27.413 | 87.734 | MN012123 | — | MN032546 |
| *N. liebigii* | JS060502_NME | Hofmann et al. 2019 | 24 | 27.407 | 87.752 | — | — | MN032547 |
| *N. liebigii* | JS060503_NME | Hofmann et al. 2019 | 24 | 27.407 | 87.752 | MN012124 | — | MN032548 |
| *N. liebigii* | KIZ-RDXZL1 | NCBI | 23 | 27.485 | 88.907 | DQ118499 | KJ810987 | HM163607 |
| *N. maculosa* | YNU-HU2002308 | NCBI | 8 | 24.400 | 100.800 | EU979835 | — | HM163588 |
| *N. medogensis* | SYNU-XZ35 | NCBI | 13 | 29.367 | 95.583 | DQ118506 | — | HM163590 |
| *N. parkeri* | N6_06_NME | Hofmann et al. 2019 | 22 | 29.589 | 90.214 | MN012125 | MN012248 | — |
| *N. parkeri* | N7_06_NME | Hofmann et al. 2019 | 22 | 29.589 | 90.214 | MN012126 | MN012249 | MN032549 |
| *N. parkeri* | N8_06_NME | Hofmann et al. 2019 | 22 | 29.589 | 90.214 | MN012127 | MN012250 | MN032550 |
| *N. parkeri* | N5_06_NME | Hofmann et al. 2019 | 21 | 29.573 | 90.433 | MN012128 | MN012251 | — |
| *N. parkeri* | TP10_06_NME | Hofmann et al. 2019 | 21 | 29.573 | 90.433 | MN012129 | MN012252 | — |
| *N. parkeri* | TP11_06_NME | Hofmann et al. 2019 | 21 | 29.573 | 90.433 | MN012130 | MN012253 | — |
| *N. parkeri* | TP8_06_NME | Hofmann et al. 2019 | 21 | 29.573 | 90.433 | — | MN012254 | — |
| *N. parkeri* | TP9_06_NME | Hofmann et al. 2019 | 21 | 29.573 | 90.433 | MN012131 | MN012255 | — |
| *N. parkeri* | N10_06_NME | Hofmann et al. 2019 | 20 | 29.578 | 90.435 | MN012132 | MN012256 | — |
| *N. parkeri* | N9_06_NME | Hofmann et al. 2019 | 20 | 29.578 | 90.435 | MN012133 | MN012257 | MN032551 |
| *N. parkeri* | TP1_06_NME | Hofmann et al. 2019 | 20 | 29.578 | 90.435 | MN012134 | MN012258 | MN032552 |
| *N. parkeri* | TP2_06_NME | Hofmann et al. 2019 | 20 | 29.578 | 90.435 | MN012135 | — | MN032553 |

| | | | | | | | | |
|---|---|---|---|---|---|---|---|---|
| *N. parkeri* | TP3_06_NME | Hofmann et al. 2019 | 20 | 29.578 | 90.435 | MN012136 | MN012259 | — |
| *N. parkeri* | CAS801L | Hofmann et al. 2019 | 19 | 30.090 | 90.480 | MN012137 | MN012260 | MN032554 |
| *N. parkeri* | CAS802L | Hofmann et al. 2019 | 19 | 30.090 | 90.480 | MN012138 | MN012261 | MN032555 |
| *N. parkeri* | CAS803L | Hofmann et al. 2019 | 19 | 30.090 | 90.480 | MN012139 | MN012262 | — |
| *N. parkeri* | CAS804L | Hofmann et al. 2019 | 19 | 30.090 | 90.480 | MN012140 | MN012263 | — |
| *N. parkeri* | CAS805L | Hofmann et al. 2019 | 19 | 30.090 | 90.480 | MN012141 | MN012264 | — |
| *N. parkeri* | A6AL_NME | Hofmann et al. 2019 | 18 | 30.156 | 90.647 | MN012142 | — | — |
| *N. parkeri* | JS0507B01_NME | Hofmann et al. 2019 | 17 | 30.378 | 90.908 | MN012143 | MN012265 | MN032556 |
| *N. parkeri* | JS0507B02_NME | Hofmann et al. 2019 | 17 | 30.378 | 90.908 | MN012144 | MN012266 | MN032557 |
| *N. parkeri* | JS0507B03_NME | Hofmann et al. 2019 | 17 | 30.378 | 90.908 | MN012145 | MN012267 | MN032558 |
| *N. parkeri* | JS0507B04_NME | Hofmann et al. 2019 | 17 | 30.378 | 90.908 | MN012146 | MN012268 | MN032559 |
| *N. parkeri* | JS0507B05_NME | Hofmann et al. 2019 | 17 | 30.378 | 90.908 | MN012147 | MN012269 | MN032560 |
| *N. parkeri* | | NCBI | 17 | 30.378 | 90.908 | KP317482 | KP317482 | HM163584 |
| *N. parkeri* | N1_06_NME | Hofmann et al. 2019 | 15 | 31.166 | 92.061 | MN012148 | MN012270 | — |
| *N. parkeri* | N2_06_NME | Hofmann et al. 2019 | 15 | 31.166 | 92.061 | MN012149 | MN012271 | — |
| *N. parkeri* | N3_06_NME | Hofmann et al. 2019 | 15 | 31.166 | 92.061 | MN012150 | MN012272 | MN032561 |
| *N. parkeri* | N4_06_NME | Hofmann et al. 2019 | 15 | 31.166 | 92.061 | MN012151 | MN012273 | — |
| *N. parkeri* | TP4_06_NME | Hofmann et al. 2019 | 15 | 31.166 | 92.061 | MN012152 | MN012274 | — |
| *N. parkeri* | TP5_06_NME | Hofmann et al. 2019 | 15 | 31.166 | 92.061 | MN012153 | MN012275 | — |
| *N. parkeri* | TP6_06_NME | Hofmann et al. 2019 | 15 | 31.166 | 92.061 | MN012154 | MN012276 | — |
| *N. parkeri* | TP7_06_NME | Hofmann et al. 2019 | 15 | 31.166 | 92.061 | MN012155 | MN012277 | — |
| *N. parkeri* | CIB-XM1096 | NCBI | 14 | 29.649 | 94.361 | DQ118498 | KJ811345 | — |
| *N. pleskei* | KQ47_14_NME | Hofmann et al. 2019 | 6 | 30.216 | 101.500 | MN012156 | MN012278 | MN032562 |
| *N. pleskei* | KQ1_14_NME | Hofmann et al. 2019 | 5 | 30.377 | 101.675 | MN012157 | MN012279 | MN032563 |
| *N. pleskei* | KQ10_14_NME | Hofmann et al. 2019 | 5 | 30.377 | 101.675 | MN012158 | MN012280 | MN032564 |
| *N. pleskei* | KQ11_14_NME | Hofmann et al. 2019 | 5 | 30.377 | 101.675 | MN012159 | MN012281 | MN032565 |
| *N. pleskei* | KQ13_14_NME | Hofmann et al. 2019 | 5 | 30.377 | 101.675 | MN012160 | MN012282 | MN032566 |
| *N. pleskei* | KQ15_14_NME | Hofmann et al. 2019 | 5 | 30.377 | 101.675 | MN012161 | MN012283 | — |
| *N. pleskei* | KQ17_14_NME | Hofmann et al. 2019 | 5 | 30.377 | 101.675 | MN012162 | MN012284 | MN032567 |
| *N. pleskei* | KQ18_14_NME | Hofmann et al. 2019 | 5 | 30.377 | 101.675 | MN012163 | MN012285 | MN032568 |
| *N. pleskei* | KQ19_14_NME | Hofmann et al. 2019 | 5 | 30.377 | 101.675 | MN012164 | MN012286 | — |
| *N. pleskei* | KQ20_14_NME | Hofmann et al. 2019 | 5 | 30.377 | 101.675 | MN012165 | MN012287 | MN032569 |
| *N. pleskei* | KQ9_14_NME | Hofmann et al. 2019 | 5 | 30.377 | 101.675 | MN012166 | MN012288 | — |
| *N. pleskei* | CAS201 | Hofmann et al. 2019 | 4 | 33.467 | 102.750 | MN012167 | MN012289 | MN032570 |
| *N. pleskei* | CAS202 | Hofmann et al. 2019 | 4 | 33.467 | 102.750 | MN012168 | MN012290 | MN032571 |
| *N. pleskei* | | NCBI | 4 | 33.467 | 102.750 | HQ324232 | HQ324232 | HM163586 |
| *N. quadranus* | SCUM20045195CJ | NCBI | 2 | 31.683 | 103.850 | DQ118514 | — | HM163591 |
| *Nanorana* sp. [A] | R5_12_NME | Hofmann et al. 2019 | 60 | 28.513 | 83.033 | MN012169 | MN012291 | MN032572 |
| *Nanorana* sp. [A] | R10_12_NME | Hofmann et al. 2019 | 59 | 28.502 | 83.129 | MN012170 | — | MN032573 |
| *Nanorana* sp. [A] | KQ2_12_NME | Hofmann et al. 2019 | 58 | 28.501 | 83.198 | MN012171 | MN012292 | MN032574 |
| *Nanorana* sp. [A] | SH070556_NME | Hofmann et al. 2019 | 53 | 28.622 | 83.662 | MN012172 | MN012293 | MN032575 |
| *Nanorana* sp. [A] | A1963/13_NME | Hofmann et al. 2019 | 52 | 28.400 | 83.700 | MN012173 | MN012294 | — |
| *Nanorana* sp. [B] | R1_09_13_NME | Hofmann et al. 2019 | 51 | 28.380 | 84.065 | MN012174 | MN012295 | MN032576 |
| *Nanorana* sp. [B] | R2_09_13_NME | Hofmann et al. 2019 | 51 | 28.380 | 84.065 | MN012175 | MN012296 | MN032577 |
| *Nanorana* sp. [B] | R4_09_13_NME | Hofmann et al. 2019 | 51 | 28.380 | 84.065 | MN012176 | MN012297 | MN032578 |
| *Nanorana* sp. [B] | SH070510_NME | Hofmann et al. 2019 | 48 | 28.074 | 85.302 | MN012177 | — | MN032579 |
| *Nanorana* sp. [C] | SH080591_NME | Hofmann et al. 2019 | 44 | 27.686 | 86.252 | MN012178 | MN012298 | MN032580 |
| *Nanorana* sp. [C] | SH080592_NME | Hofmann et al. 2019 | 44 | 27.686 | 86.252 | MN012179 | MN012299 | MN032581 |
| *Nanorana* sp. [C] | SH080593_NME | Hofmann et al. 2019 | 44 | 27.686 | 86.252 | MN012180 | MN012300 | MN032582 |
| *Nanorana* sp. [C] | SH080594_NME | Hofmann et al. 2019 | 44 | 27.686 | 86.252 | MN012181 | MN012301 | MN032583 |

| | | | | | | | | |
|---|---|---|---|---|---|---|---|---|
| *Nanorana* sp. [C] | SH080570_NME | Hofmann et al. 2019 | 43 | 27.697 | 86.275 | MN012182 | MN012302 | MN032584 |
| *Nanorana* sp. [C] | SH080571_NME | Hofmann et al. 2019 | 43 | 27.697 | 86.275 | MN012183 | MN012303 | MN032585 |
| *Nanorana* sp. [C] | SH080572_NME | Hofmann et al. 2019 | 43 | 27.697 | 86.275 | MN012184 | MN012304 | MN032586 |
| *Nanorana* sp. [C] | SH080553_NME | Hofmann et al. 2019 | 41 | 27.718 | 86.311 | MN012185 | MN012305 | MN032587 |
| *Nanorana* sp. [C] | SH080555_NME | Hofmann et al. 2019 | 41 | 27.718 | 86.311 | MN012186 | MN012306 | MN032588 |
| *Nanorana* sp. [C] | SH080545_NME | Hofmann et al. 2019 | 40 | 27.703 | 86.337 | MN012187 | MN012307 | MN032589 |
| *Nanorana* sp. [C] | SH080546_NME | Hofmann et al. 2019 | 40 | 27.703 | 86.337 | MN012188 | MN012308 | MN032590 |
| *Nanorana* sp. [C] | SH080548_NME | Hofmann et al. 2019 | 40 | 27.703 | 86.337 | MN012189 | MN012309 | MN032591 |
| *Nanorana* sp. [C] | SH080551_NME | Hofmann et al. 2019 | 40 | 27.703 | 86.337 | MN012190 | MN012310 | MN032592 |
| *Nanorana* sp. [C] | SH080552_NME | Hofmann et al. 2019 | 40 | 27.703 | 86.337 | MN012191 | MN012311 | — |
| *Nanorana* sp. [C] | SH080512_NME | Hofmann et al. 2019 | 39 | 27.595 | 86.340 | MN012192 | MN012312 | MN032593 |
| *Nanorana* sp. [C] | SH080523_NME | Hofmann et al. 2019 | 37 | 27.694 | 86.351 | MN012193 | MN012313 | MN032594 |
| *Nanorana* sp. [C] | Ne13_13_NME | Hofmann et al. 2019 | 35 | 27.576 | 86.514 | MN012194 | MN012314 | MN032595 |
| *Nanorana* sp. [C] | Ne1_13_NME | Hofmann et al. 2019 | 32 | 27.689 | 86.731 | MN012195 | MN012315 | MN032596 |
| *Nanorana* sp. [C] | Ne2_13_NME | Hofmann et al. 2019 | 32 | 27.689 | 86.731 | MN012196 | MN012316 | MN032597 |
| *Nanorana* sp. [C] | Ne9_13_NME | Hofmann et al. 2019 | 31 | 27.671 | 86.765 | MN012197 | MN012317 | MN032598 |
| *Nanorana* sp. [Chainpur] | A1966/13_NME | Hofmann et al. 2019 | 67 | 29.374 | 81.137 | MN012198 | MN012318 | MN032599 |
| *N. vicina* | 2Bhan_RAS | Hofmann et al. 2019 | 74 | 32.873 | 75.858 | MN012199 | MN012319 | — |
| *N. vicina* | 1G_RAS | Hofmann et al. 2019 | 73 | 32.777 | 75.947 | — | MN012320 | MN032600 |
| *N. vicina* | 1Pa_RAS § | Hofmann et al. 2019 | 72 | 32.528 | 75.991 | — | MN012321 | MN032601 |
| *N. vicina* | 2Ba_RAS | Hofmann et al. 2019 | 71 | 31.783 | 77.068 | — | — | MN032602 |
| *N. vicina* | 2Baj_RAS | Hofmann et al. 2019 | 70 | 31.821 | 77.112 | — | — | MN032603 |
| *N. vicina* | 2Pul_RAS § | Hofmann et al. 2019 | 69 | 31.996 | 77.448 | MN012200 | — | MN032604 |
| *N. vicina* | 782_RAS § | Hofmann et al. 2019 | 68 | 31.261 | 77.450 | MN012201 | — | MN032605 |
| *N. taihangnica* | | NCBI | 1 | 35.265 | 112.090 | KF199146 | KF199146 | HM163608 |
| *N. unculuanus* | YNUHU2002502601 | NCBI | 7 | 24.447 | 100.834 | DQ118491 | — | HM163595 |
| *N. ventripunctata* | SCUM045887WD | NCBI | 11 | 27.830 | 99.701 | EU979839 | KJ810985 | HM163585 |
| *N. ventripunctata* | SH050538_NME | Hofmann et al. 2019 | 10 | 27.788 | 99.855 | MN012208 | MN012328 | MN032610 |
| *N. ventripunctata* | SH050539_NME | Hofmann et al. 2019 | 10 | 27.788 | 99.855 | MN012209 | MN012329 | MN032611 |
| *N. yunnanensis* | | NCBI | 9 | 27.724 | 100.789 | KF199150 | KF199150 | HM163593 |
| *Q. boulengeri* | YNU-HU20025106 | NCBI | 80 | 28.811 | 105.831 | KX645665 | KX645665 | HM163604 |
| *Q. delacouri* | FMNH255623 | NCBI | 82 | 19.018 | 104.799 | EU979810 | EU979664 | HM163600 |
| *Q. exilispinosa* | KF199151 | NCBI | 81* | 22.396 | 114.109 | KF199151 | KF199151 | HM163610 |
| *Q. jiulongensis* | KF199149 | NCBI | 75* | 27.750 | 117.683 | KF199149 | KF199149 | HM163603 |
| *Q. shini* | KF199148 | NCBI | 76 | 25.598 | 109.935 | KF199148 | KF199148 | HM163602 |
| *Q. spinosa* | | NCBI | 77 | 24.481 | 99.047 | NC_013270 | NC_013270 | HM163606 |
| *Q. verrucospinosa* | | NCBI | 78 | 21.789 | 101.142 | KF199147 | KF199147 | HM163599 |
| *Q. yei* | HNNU0908I061 | NCBI | 79* | 31.798 | 115.407 | KJ842105 | KJ842105 | HM163596 |
| *Fejervarya cancrivora* | | NCBI | | | | EU652694 | EU652694 | HM163581 |
| *Hoplobatrachus* [illegible] | | NCBI | | | | NC_019615 | NC_019615 | HM163612 |
| *Limnonectes fragilis* | ZNAC11006 | NCBI | | | | AY899241 | AY899241 | HM163611 |
